# Multi-omics approach identifies novel pathogen-derived prognostic biomarkers in patients with *Pseudomonas aeruginosa* bloodstream infection

**DOI:** 10.1101/309898

**Authors:** Matthias Willmann, Stephan Götting, Daniela Bezdan, Boris Maček, Ana Velic, Matthias Marschal, Wichard Vogel, Ingo Flesch, Uwe Markert, Annika Schmidt, Pierre Kübler, Maria Haug, Mumina Javed, Benedikt Jentzsch, Philipp Oberhettinger, Monika Schütz, Erwin Bohn, Michael Sonnabend, Kristina Klein, Ingo B Autenrieth, Stephan Ossowski, Sandra Schwarz, Silke Peter

**Author notes:** Address correspondence to Matthias Willmann: Institute of Medical Microbiology and Hygiene, Elfriede-Aulhorn-Str 6, 72076, Tübingen, Germany.

## Abstract

*Pseudomonas aeruginosa* is a human pathogen that causes health-care associated blood stream infections (BSI). Although *P. aeruginosa* BSI are associated with high mortality rates, the clinical relevance of pathogen-derived prognostic biomarker to identify patients at risk for unfavorable outcome remains largely unexplored. We found novel pathogen-derived prognostic biomarker candidates by applying a multi-omics approach on a multicenter sepsis patient cohort. Multi-level Cox regression was used to investigate the relation between patient characteristics and pathogen features (2298 accessory genes, 1078 core protein levels, 107 parsimony-informative variations in reported virulence factors) with 30-day mortality. Our analysis revealed that presence of the *helP* gene encoding a putative DEAD-box helicase was independently associated with a fatal outcome (hazard ratio 2.01, p = 0.05). *helP* is located within a region related to the pathogenicity island PAPI-1 in close proximity to a *pil* gene cluster, which has been associated with horizontal gene transfer. Besides *helP*, elevated protein levels of the bacterial flagellum protein FliL (hazard ratio 3.44, p < 0.001) and of a bacterioferritin-like protein (hazard ratio 1.74, p = 0.003) increased the risk of death, while high protein levels of a putative aminotransferase were associated with an improved outcome (hazard ratio 0.12, p < 0.001). The prognostic potential of biomarker candidates and clinical factors was confirmed with different machine learning approaches using training and hold-out datasets. The *helP* genotype appeared the most attractive biomarker for clinical risk stratification due to its relevant predictive power and ease of detection.

## Introduction

Blood stream infections (BSI) are a frequent and often fatal occurrence in hospitalized patients, particularly under immunosuppression (ECDC 2015). According to the European Detailed Mortality Database (http://data.euro.who.int/dmdb/), more than 40,000 deaths in Europe can be attributed to sepsis in 2014. *Pseudomonas aeruginosa* is an important pathogen causing up to 15.4% of all BSI cases (ECDC 2015). Mortality rates of up to 42% even in advanced settings are reported (McCarthy and Paterson 2017), especially when appropriate antibiotic treatment is delayed (Skaar 2010).

The search for appropriate biomarkers is linked with the prospect of improving early diagnosis and prognosis prediction in sepsis. To date, C-reactive protein, interleukin-6 and procalcitonin are the only well-established diagnostic biomarkers, despite extensive evaluation of more than 100 biomarkers (Pierrakos and Vincent 2010). The majority of these biomarkers is host-derived. This is in line with the current paradigm of sepsis pathophysiology that explains lethal septic shock and multi-organ failure primarily as a result of the host’s pro- and anti-inflammatory reaction to pathogen components like carbohydrate and fatty acids, termed pathogen-associated molecular patterns (PAMP) (Walton et al. 2014; Gotts and Matthay 2016). It is indeed well known that the genetic diversity in human genes encoding for pathogen recognition receptors as well as for pro- and anti-inflammatory mediators explains in part the variability in the clinical course of septic patients (Khor et al. 2007; Lehmann et al. 2009; Thompson et al. 2014). However, the role of the nature and characteristics of the infecting pathogen is frequently neglected (Lisboa et al. 2010; Angus and van der Poll 2013). Given the huge diversity of bacterial genomes and functional capacities even within one species, the pathogen itself could account for unexplained heterogeneity in the clinical course and outcome of sepsis. Recently, the pangenome of the species *P. aeruginosa* was estimated to contain more than 16,000 non-redundant genes, while only 15% of these genes were present in all strains forming the core genome (Mosquera-Rendon et al. 2016). Of particular interest are prognostic bacterial biomarkers that can indicate the risk of a fatal outcome in septic patients, thus providing guidance in therapy and improved management of patient monitoring.

While bacterial virulence factors have been extensively explored in *P. aeruginosa* (Veesenmeyer et al. 2009), investigations have almost exclusively been carried out in *in vitro* experimentations or in animal models providing no evidence of their relevance and utility as prognostic biomarkers in humans. The type 3 secretion bacterial effector proteins ExoS, ExoT, and ExoU are an exception (Lisboa et al. 2010), with some authors reporting an association between expression level and sepsis outcome (El-Solh et al. 2012; Hattemer et al. 2013). In addition, one recent study presented evidence that the presence of the *exoU* gene is an independent predictor of early sepsis mortality in *P. aeruginosa* BSI (Pena et al. 2015).

In a multicenter study, we applied a multi-omics approach to identify pathogen factors that contribute to differential mortality outcomes in patients with *P. aeruginosa* bloodstream infection. We first used genomics and proteomics to characterize *P. aeruginosa* strains from sepsis patients. Next, we integrated these omics data from bacterial isolates with treatment- and patient-related data to gain a broader understanding of the complex interactions between host and pathogen during blood stream infections. We thereby screened for pathogen factors which are independently linked to 30-day mortality and which would consequently be attractive prognostic biomarker candidates. Finally, we confirmed biomarker candidates identified by our statistical model using different machine learning algorithms.

## Results

### Clinical and patient-related risk factors for 30-day mortality

From 175 eligible patients, 166 (94.86%) patients with *P. aeruginosa* BSI were included into the final analysis (Figure S1). The basic demographic, clinical and infection-related characteristics of the patient study population are presented in table S1. An investigation of factors that had an impact on the mortality rate was initially performed on clinical and patient-related variables (Table S2). Multivariate Cox regression modelling revealed that immunosuppression as well as a rise in the SAPS II score increased 30-day mortality while administration of appropriate antibiotic treatment and a genitourinary infection source decreased the risk of a fatal outcome (Table S3).

### Genomic characteristics of clinical *P. aeruginosa* strains

The genome of the first isolate recovered from each patient with a *P. aeruginosa* blood stream infection was sequenced (166 strains). The pangenome consisted of 23917 genes with 4354 core genes shared by > 99% of isolates, 639 soft core genes shared by 95% - 99%, 1762 shell genes shared by 15% - 95%, and 17162 cloud genes shared by < 15%. The high number of accessory genes underlines the stupendous genomic diversity and plasticity of *P. aeruginosa* species represented by our study dataset.

The phylogenetic tree based on the core genome SNP alignment shows a highly diverse population structure of our clinical isolates with distinct clades of similar branch length within two major phylogenetic clusters along with a discriminative cluster formed by only 3 strains (Figure 1). Recombination had occurred at a median rate of 0.07 (SNPs inside recombination/SNPs outside recombination, interquartile range: 0.03 - 0.27), demonstrating that recombination events have only slightly contributed to shaping the diversity of our strain set. Closely related isolates were most commonly found in only one hospital in a narrow time frame, providing evidence for a spatial and temporal clustering. In some cases, strains from the same cluster have been isolated in different hospitals during the entire study period, suggesting either a transfer from one to the other hospital or a circulation of the particular strain within the community and sporadic reintroduction in our hospitals.

**Figure 1.**
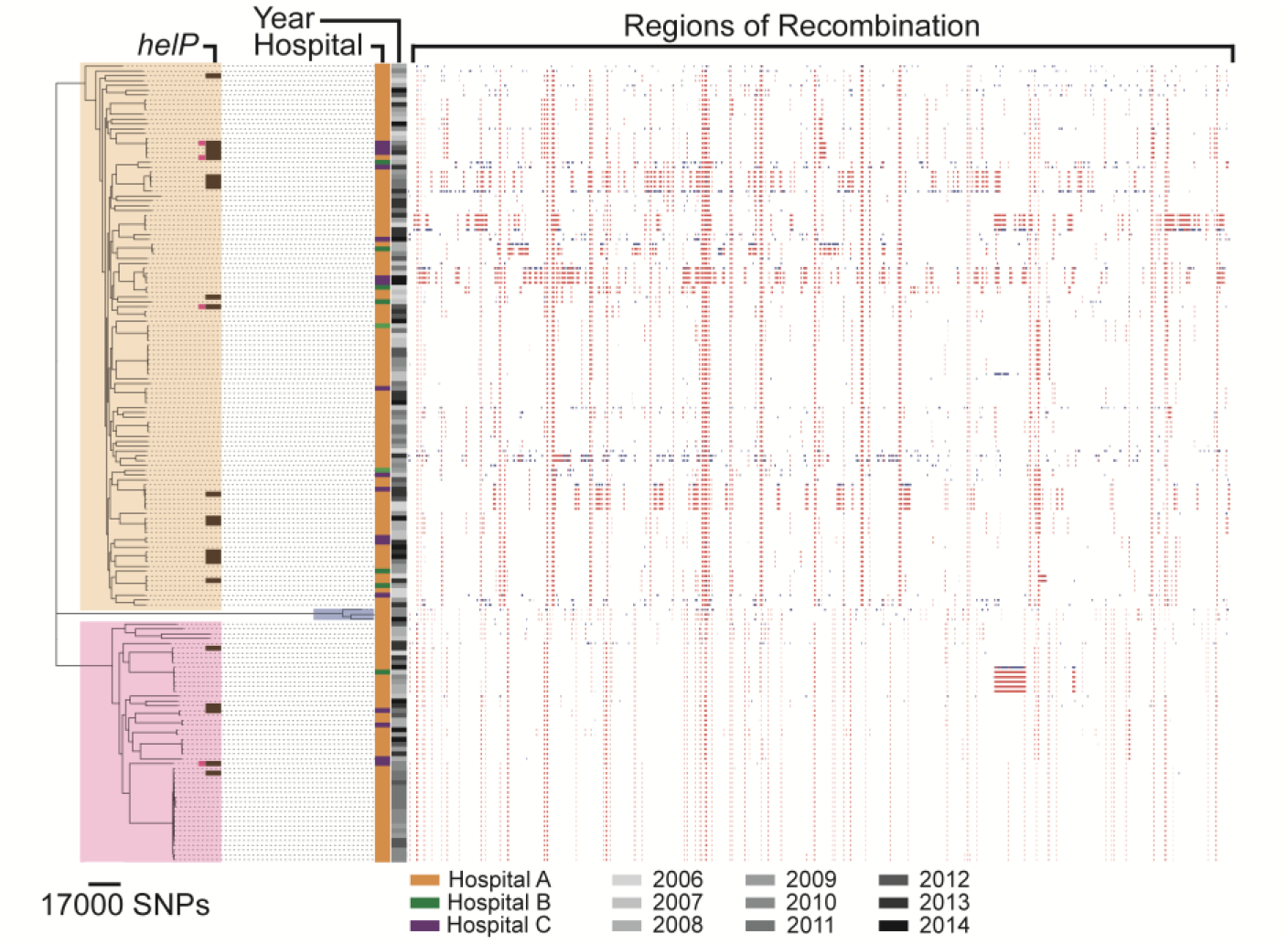
Core genome phylogenetic, temporal and spatial relationship of clinical *P. aeruginosa* strains recovered from patients with blood stream infection. A maximum-likelihood phylogeny of the core genome SNP alignment reveals three major genomic clusters (Core-cluster 1 = bright yellow; Core-cluster 2 = light rose; Core-cluster 3 = grey). A high diversity within these clusters is reflected by numerous subgroups and distinct single isolates. Location and year of isolation is provided for each strain. Regions of predicted recombinations are shown by the right-sided panels of blocks. Red blocks indicate recombinations on internal branches, therefore shared by several strains through common descent. Blue blocks indicate recombinations that take place on terminal branches, thus are specific to individual isolates. Presence of the dead box helicase gene *helP* is shown by brown blocks beside the phylogenetic tree, with pink-colored squares that illustrate strains where *helP* location was predicted to be on a plasmid (plasmidSPAdes). The scale beneath demonstrates a distance of 17.000 point mutations.

Antibiotic susceptibility profiles are shown in figure S2, demonstrating a wide range from broadly susceptible to extensively drug resistant (XDR) strains. XDR strains were only susceptible to colistin and were usually phylogenetically clustered, suggesting outbreak situations as previously described (Willmann et al. 2015).

In order to investigate the genomic relatedness of the accessory genome between the 166 strains, we only considered accessory genes with a prevalence ≥ 10% and ≤ 90%. Using this criterion, a subset of 2298 accessory genes was tested. Ward analysis revealed that the 166 isolates can be divided in four accessory genome (acc-) clusters (Figure 2A). Except for acc-cluster 4, which was confined to a 4-year period, the appearance of strains from the other acc-clusters was evenly distributed over time (Figure 2B). Acc-clusters showed a strong affiliation to the three major core-genome clusters, underlining a further structural distinction within the core genome phylogeny (Figure 2C). Acc-clusters were included in the clinical Cox regression model. The analysis showed that acc-cluster 2 was independently associated with 30-day mortality (HR 1.95, p = 0.048, Wald test), suggesting the presence of genomic pathogen factors that negatively influence patient survival (Table S4).

**Figure 2.**
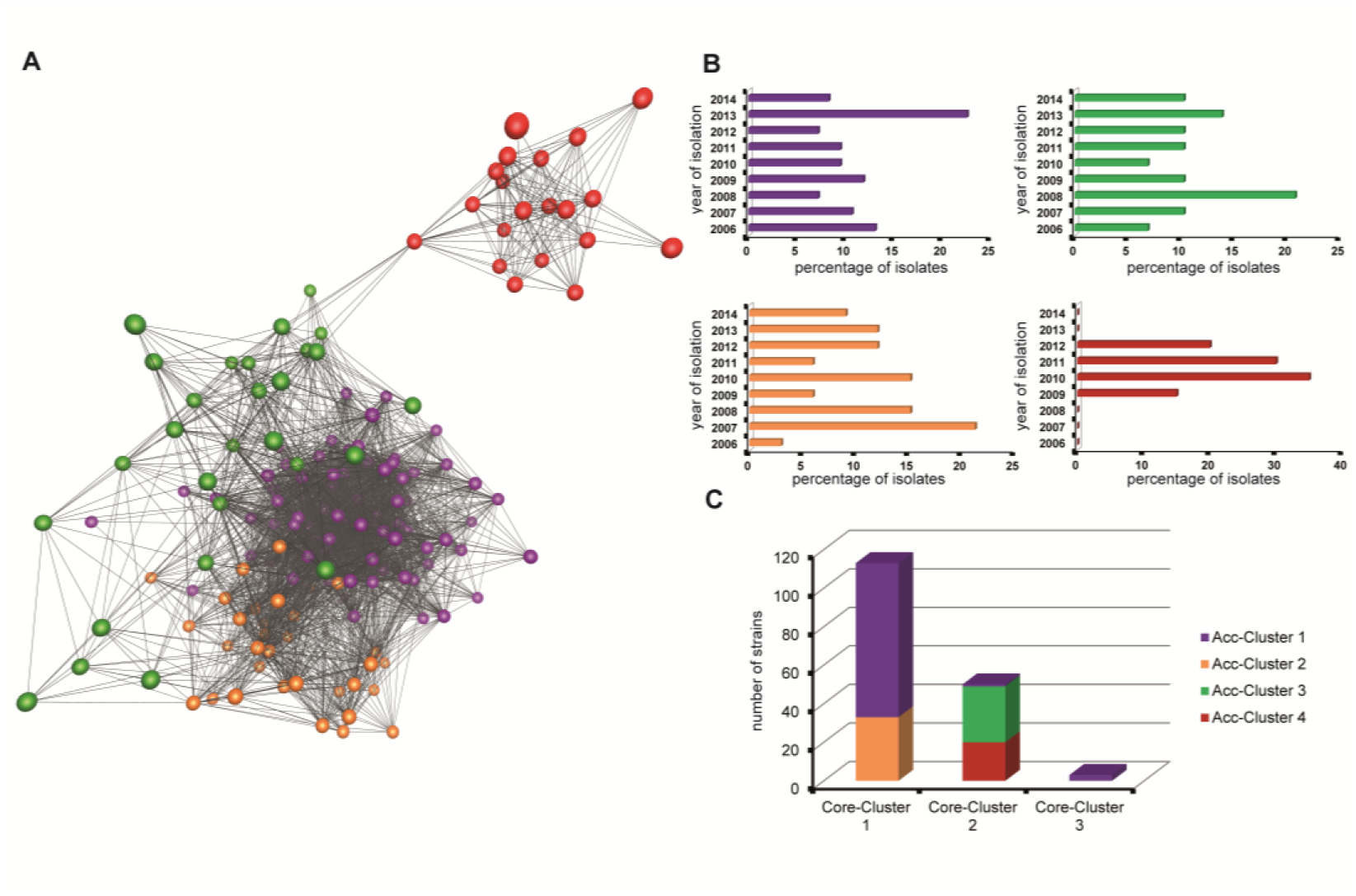
Genomic and temporal clustering of accessory genes and its link to core genome clusters. (A) Population structure of the accessory genome using the Ward cluster algorithm with the Pearson similarity index and displayed as three-dimensional correlation network by Biolayout. Each node represents a study isolate, whereas the connecting edges reflect similarity based on the input gene presence-absence matrix. Four accessory genome clusters (acc-clusters) were revealed, coded by different colors (acc-cluster 1 = purple; acc-cluster 2 = orange; acc-cluster 3 = green, acc-cluster 4 = red). (B) The histograms illustrate the distribution of isolation time over the study period for each acc-cluster. All acc-clusters were evenly isolated with the exception of acc-cluster 4 which has only been found consecutively in four years (2009 - 2012). (C) The bars display overlaps between core- and acc-clusters. Core-cluster 1 and 2 were mainly partitioned in two distinct types of accessory genomes, hence rising evidence for a deeper structural genomic disparity than the one shown by the core genome maximum-likelihood phylogeny. Only one isolate from core-cluster 2 had an accessory genome that is grouped with acc-cluster 1. The three isolates of core-cluster 3 belong exclusively to acc-cluster 1, pointing to a closer genomic relation to core-cluster 1 than core cluster 2.

Subsequently, we investigated whether certain gene ontology (GO) terms and thereby gene functions are over- or under-represented in acc-cluster 2. Compared to the three remaining acc-clusters that served as a reference, acc-cluster 2 was enriched with the GO terms “peptidyl-histidine modification” (GO:0118202, FDR = 0.033) and “peptidyl-histidine phosphorylation” (GO:0018106, FDR = 0.033) (Figure S3). Both GO terms involve sensor histidine kinase genes that usually function in two-component systems. These bacterial regulatory systems, designed to sense external stimuli and to facilitate an appropriate adaptive response to stressors and changes in environmental and growth conditions, modulate the transcription of genes including virulence factors and antimicrobial resistance genes in *P. aeruginosa* (Gooderham and Hancock 2009). Such systems could have a significant influence on a strain’s survival chance during infection (Mikkelsen et al. 2011).

### Protein level characteristics of clinical *P. aeruginosa* strains

After determining the genomic features, we next defined the cellular proteome of all 166 isolates. A total of 7757 unique proteins were identified in the proteomics analysis, with a subset of 1078 proteins (13.9%) synthesized by all study strains (core proteome). Principal component analysis of protein level profiles of the strains did not show clustering according to the survival status of patients, neither for the core-proteome (Figure 3A) nor for the whole proteome (Figure 3B). Ward cluster analysis of the core proteome resulted in four core proteome clusters (prot-clusters) of strains with closely related protein level profiles (Figure 3C). All prot-clusters were found in both the two numerically greatest core genome clusters (Figure 3D). This demonstrates that even phylogenetically distinct strains can share similar profiles of core protein levels. Since the presence of different accessory genes could also have an impact on the protein level pattern, we ordered the strains according to their acc-clusters and compared the protein levels of the core proteome (Figure S4A). We observed no distinct patterns associated with an acc-cluster or a strong relationship in a cross-comparison of prot- and acc-clusters (Figure S4B).

**Figure 3.**
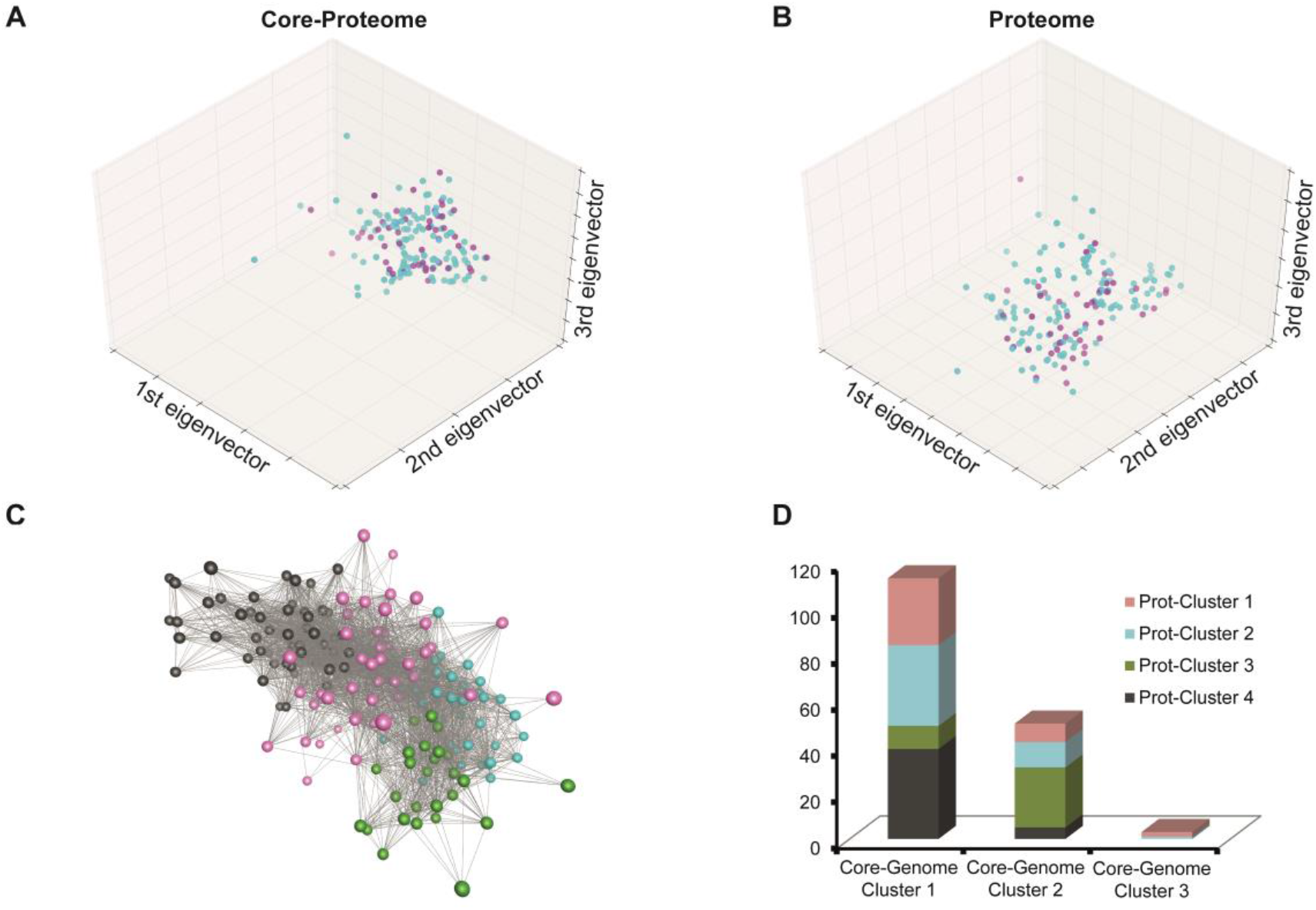
Basic proteome and core proteome characteristics and comparison with genomic features. A principle component analysis with the first three eigenvectors is presented for the core proteome (A) and the whole proteome (B). Any data point stands for a strain isolated from survivors (blue) and decedents (purple). In both cases, no clustering according to survival status was observed. The first three eigenvectors comprised 57.1% of the overall variance for the core proteome, and 68.51% for the proteome. (C) Four core-proteome clusters were revealed, each of them representing a highly correlated expression profile. Visualization was done using Biolayout. Clusters are labelled as indicated by the color bar in D. (D) The fraction of the four identified core proteome clusters within the three core genome clusters showed the presence of all closely related protein level profiles in both the numerical greatest core genome clusters. This suggested that the protein level profiles are independent from the underlying core genome structure.

Inclusion of prot-clusters into the clinical Cox regression model did not reveal a link between prot-clusters and 30-day mortality (Table S4). These results suggest that the risk of a fatal outcome is not determined by complex core proteome clusters. But since individual protein levels could still have an impact, we performed a multi-level Cox regression analysis of single genomic and protein level factors on patient outcome.

### Multi-level Cox regression and prediction model

Figure S5 provides an overview of the statistical workflow up to the final prediction model using a multi-level Cox regression analysis approach. We conducted an in-depth analysis of a pre-assigned accessory genome subset comprising 2298 genes and the natural log-normalized protein level data set of 1078 core proteins. Four pathogen-derived factors (one genomic and three proteomic factors) were independently associated with mortality in the final Cox regression prediction model (Table 1).

**Table 1.**
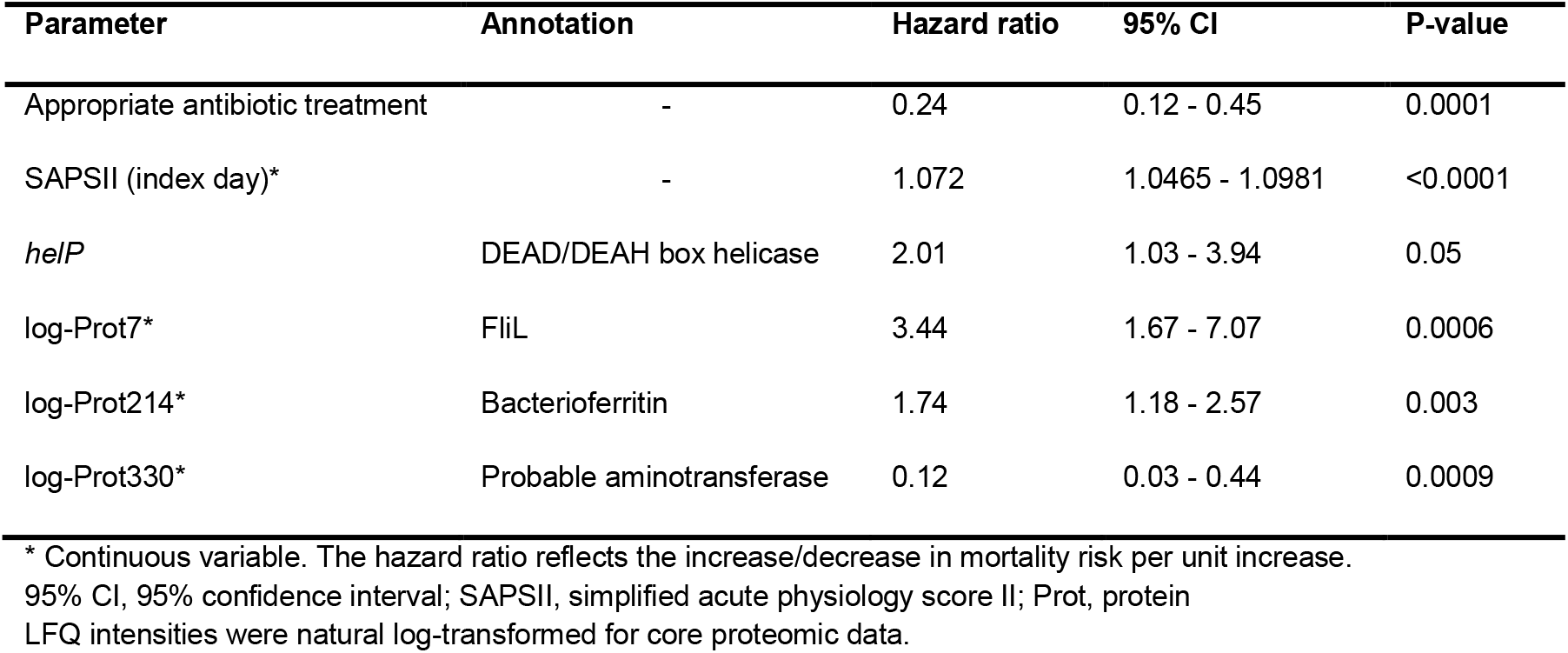
Final multivariate Cox regression model including virulence factor candidates

Prior screening model values of the four pathogen-derived predictors are presented in table S5, while all factors that were included into the multivariate models are shown in table S6. Two variables of the clinical Cox regression model were removed from the final and comprehensive Cox regression prediction model: immunosuppression due to a p-value > 0.05 and genitourinary infection source due to failing the internal bootstrap validation process. Three of the four pathogen-derived predictors shown in table 1 increase the risk of death (hazard ratio > 1), either when present in the bacterial genome (the genomic factor *helP*) or in case of high protein levels (proteins Prot7 and Prot214). All three were considered pathogen-derived risk factors for a fatal outcome.

In contrast, high levels of Prot330, a putative aminotransferase, turned out to have an anti-virulence effect (hazard ratio 0.12, 95% CI 0.03 - 0.44, p = 0.0023). Table S7 illustrates the functional annotation of genomic and proteomic predictors and their GO-terms from the UniProtKB database (http://www.uniprot.org/).

Besides protein levels and presence of genes, the existence of structural variations in putative pathogen virulence factors within the core genome set could have further contributed to mortality and was thus specifically explored. A total of 92 reported putative virulence factors were identified in the dataset; 59 of those were present in the core genome (Table S8). Using PAO1 (accession number: NC_002516.2) and PA14 (accession number: NC_008463.1) as genetic references, multivariate analyses of 107 parsimony-informative SNPs causing amino acid replacements detected only one candidate (LasA, A111V) linked to mortality (hazard ratio 2.18; 95% confidence interval 1.16 - 4.09; p = 0.012) in the variant screening model. However, this SNP candidate failed statistical significance in the final Cox regression prediction model (p = 0.28), suggesting that structural variations in putative pathogen virulence factors did not impact 30-day mortality in our study population.

### In-depth characterization of the prognostic biomarker candidate *helP*

One of the identified prognostic biomarker candidates, the 1866^th^ accessory genome gene that we named *helP* (GenBank accession number: KY940721), is particularly interesting. The presence of *helP* in the genome of the study strains was estimated to double the risk of a fatal outcome (hazard ratio 2.01, 95% confidence interval 1.03 - 3.94, p = 0.05). Its gene product is highly similar to RL063, a protein whose gene sequence is located on the pathogenicity island I in PA14 (98.3% protein sequence similarity, UniProt accession number Q7WXZ7). The amino acid sequence of HelP is identical to a protein named PSPA7_4493 (UniProt accession number: A6V9V7), a predicted DEAD/DEAH box helicase from the *P. aeruginosa* strain PA7. This prediction is primarily based on a helicase conserved C-terminal domain (PF00271, domain boundary positions: 570 - 685) and a DEXDc domain (SM00487, domain boundary positions: 57 - 410). *helP* appeared in 22 of our study strains, but much more frequently in the high-risk acc-cluster 2 strains (27.27% in acc2 vs 9.77% in the other three clusters, p = 0.008, chi-squared test). A maximum-likelihood phylogeny showed that HelP groups together with other predicted helicases from Pseudomonas sp., thereby most closely related to the class of DEAD-box helicases within the superfamily 2 (Figure S6, table S9).

Generally, *helP* was evenly distributed among all strains in the core phylogeny and was found in different hospitals (Figure 1), reflecting the gene’s integration in many different phylogenetic groups rather than just in one. We hypothesized that *helP* might be transmittable via horizontal gene transfer, which is another important aspect apart from virulence capabilities. DEAD/DEAH box helicases are non-essential bacterial genes that might be acquired through horizontal gene transfer. The recycler tool (Rozov et al. 2016) and plasmidSPAdes (Antipov et al. 2016) were used to predict plasmids from the short Illumina sequence reads of all *helP* positive strains. Plasmids predicted by plasmidSPAdes harbored *helP* in strain ID 26, ID 93, ID 101, and ID 138 (Figure 1) while plasmids predicted by the recycler tool did not. In the remaining strains, *helP* location was predicted to be on the bacterial chromosome. Since genome assembly from short reads can be prone to errors, particularly in the detection and characterization of mobile genetic elements, we sequenced the four strains including strain ID50 on a PacBio instrument to improve assembly quality. In all strains, *helP* was located on a contig with a size > 800 kb. This makes a location on a plasmid very unlikely, indicating that plasmidSPAdes provided a false positive rating.

Nevertheless, *helP* can be found in 12 strains that originate from four different phylogentic clusters as well as in 10 strains that are genetically distinct from all other *helP* positive strains. This genomic diversity of *helP* positive strains suggests a horizontal transfer of *helP* in the past. Interestingly, the genomic environment of *helP* on the five large PacBio contigs resembled the pathogenicity island PAPI-1 from PA14 (Figure S7), where a homologous gene of *helP* is located (RL063). Particularly upstream of *helP*, we found PAPI-1 well conserved. Of special interest is the 10-gene cluster of a type IV pilus (T4P) apparatus that is located in close proximity to *helP* (Figure S7). This T4P system has been described as a conjugative apparatus genetically closely related to genes on the enterobacterial plasmid R64. The system has been reported of being capable of transferring PAPI-1 into recipient *P. aeruginosa* (Carter et al. 2010). When mapping the short Illumina sequence reads of the 22 *helP* positive strains against PAPI-1, we found a similar picture with a few differences between single strains and clusters (Figure S8). The structure of this genomic environment suggests a past exchange of *helP* between different *P. aeruginosa* strains via conjugation machineries but not via plasmids.

It was recently reported that a RNA helicase (Uniprot accession: Q9I003) in *P. aeruginosa* affected expression of ExoS (Tan et al. 2016). For this reason, we explored protein levels of known *P. aeruginosa* exotoxins, secretion system effectors and factors of the T4P in all clinical isolates (Table S10). Strains that were positive for *helP* had a 6.57-fold higher expression of *exoU* compared to *helP* negative strains (p = 0.04), but were not distinct in their ExoS levels. This is likely due to different structures of both helicases. In a pairwise alignment comparison, HelP had only a 20% amino acid sequence similarity with the respective RNA helicase (Uniprot accession: Q9I003), suggesting that both putative helicases do not necessarily operate with the same mode of action. However, our findings indicate a potential connection between the *helP* genotype and ExoU expression, and could play a role in sepsis when considering the clinical impact of the *exoU* genotype (Pena et al. 2015).

### Predictive performance of identified prognostic factors in machine learning algorithms

The following datasets were submitted for further evaluation using different machine learning strategies: datasets including all features of the three screening models (genomic, phenotypic, or SNP features), a dataset containing all pathogen-derived factors from the screening models (“ALL”) and the dataset with the variables from the final Cox regression model (“Final”). All datasets consisted of the clinical predictors identified in the clinical Cox regression model (Table S3).

Performance specifications of the estimators from each tested dataset are presented in supplement table S11. Values for the area under the receiver operating characteristic curve (AUC) indicate each estimator’s potential to discriminate patients at high risk of a fatal outcome from those with a lower risk. In most cases, AUC values were below 0.8, indicating a rather weak discriminatory power of the estimators. Exceptions were estimators from the dataset of the final cox regression model which gained higher AUC values compared with the estimators from the other datasets (median AUC 0.829 vs 0.736, p = 0.0009). The best estimator from the dataset of the final cox regression model was a linear support vector classifier that showed no sign of overfitting in its learning curve (Figure 4A). In contrast, most estimators were prone to overfitting and were difficult to regularize. However, using the best 5% of features generally increased performance significantly and often removed overfitting. This indicates a high background of uninformative features that disturb the predictive potential of the estimators. It underlines the importance of feature selection in datasets with a high number of features and a comparative smaller number of instances, as is usually the case in multi-omics studies.

**Figure 4.**
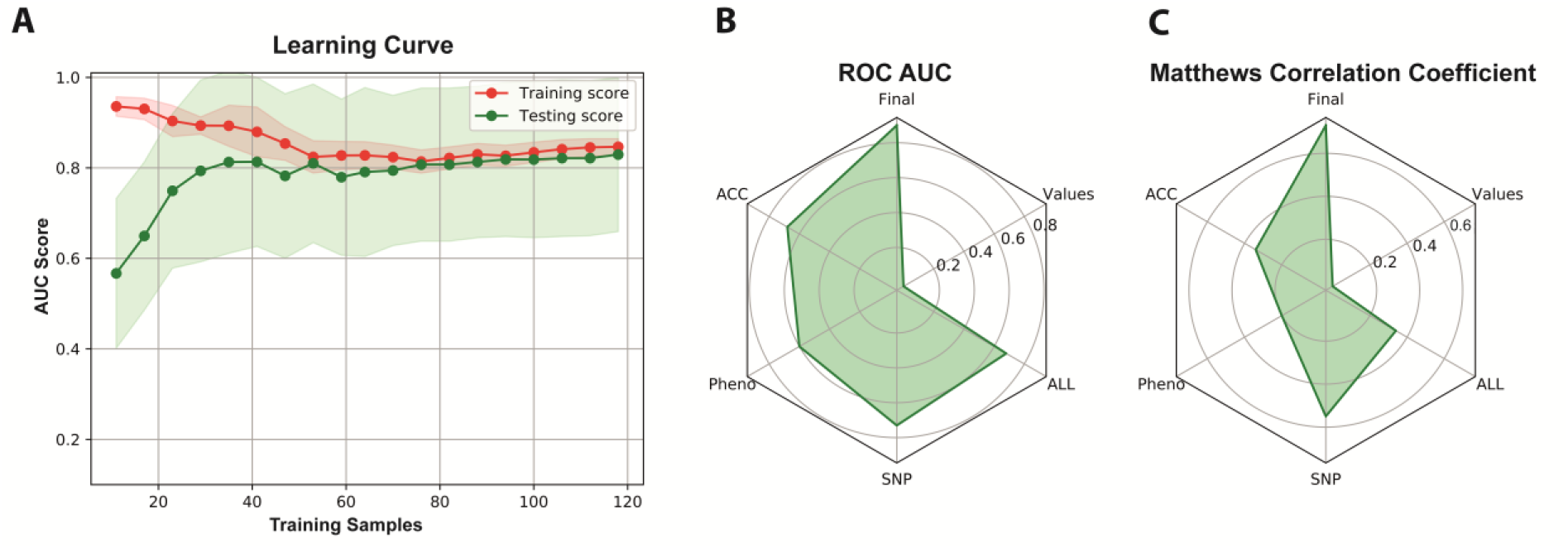
Machine learning estimator validation of the multi-omics datasets. A) The learning curve of the linear support vector machine estimator from the dataset containing the features from the final Cox regression model (“Final”) determined cross-validated training and test area under the receiver operating characteristic curves scores (AUC scores) for different training set sizes. The faded areas indicate standard deviations of the respective test scores. Both curves approach each other with increasing training set size, reflecting evidence against overfitting. (B) The radial graph demonstrates the area under the receiver operating characteristic curve (ROC AUC) values of the final validation step, when the best estimators from each dataset were tested against the hold-out data. The prediction performance from estimators of the following datasets was examined: ACC = accessory genome features, Pheno = protein levels and antibiotic susceptibility features, SNP = single nucleotide polymorphisms in reported virulence factors, ALL = combination of the three models above. These datasets included the clinical risk factors from the respective clinical Cox regression model. “Final” marks the estimator that consists of the dataset with the features from the final Cox regression model. The label “Values” does not indicate a dataset but the AUC values at the grid in the radial graph. The estimator from the Final dataset showed superior performance (AUC = 0.895), particularly over the Pheno dataset estimator which performed quite weak (AUC = 0.595). (C) Matthews correlation coefficients are provided for the same estimators as mentioned in B. The label “Values” indicates the correlation coefficients but not a dataset. Again, the estimator from the Final dataset shows the highest coefficient (0.726), reflecting a high number of correct predictions.

For each dataset, we subsequently evaluated the most promising estimator on the hold-out dataset that contained 20% of the total data (Table S12). The estimator trained on the dataset with the features from the final Cox regression model was clearly superior to the estimators of the other datasets (AUC = 0.895, Matthews correlation coefficient 0.726, Figure 4B and 4C). It was the only model that predicted all fatal cases correctly and thus did not miss one patient at high risk of a fatal outcome (Table S12). This suggests a significant relation between the identified clinical risk factors and prognostic biomarker candidates with fatal outcome. It also demonstrates that a feature selection based on classical epidemiological methodology (here Cox regression) can be a powerful tool when combined with machine learning algorithms in multi-omics approaches.

## Discussion

Our approach integrated genomic and proteomic pathogen data into a clinical multicenter cohort study with a wide range of sepsis conditions. This was followed by an evaluation of promising prognostic biomarker candidates using machine learning. Our results support that certain *P. aeruginosa* pathogen factors significantly contribute to the risk of a fatal outcome in bloodstream infections independently of the physiological patient status and administration of appropriate antibiotic therapy.

In terms of a more detailed characterization, we have focused on the genomic candidate *helP* since its presence increased the risk of death by two-fold in our study and since it can be easily measured in a routine diagnostic setting (e.g. by PCR), which makes it an interesting prognostic biomarker. Moreover, its genomic environment and its detection in different genetic lineages suggest that *helP* has been acquired by horizontal gene transfer.

We found *helP* in close genomic proximity to a type IV pili (T4P) gene complex within a genetic environment that is similar to the pathogenicity island I of PA14. These T4P systems are involved in motility and adhesion to host cells during infection (Hahn 1997; Bieber et al. 1998). T4P-deficient *P. aeruginosa* mutants were reported to have a lower cytotoxicity, potentially due to the loss of cell contact and therefore an inefficiently working type III secretion system (T3SS) (Comolli et al. 1999), whose importance as a prognostic biomarker in *P. aeruginosa* bacteremia has been repeatedly shown (El-Solh et al. 2012; Hattemer et al. 2013; Pena et al. 2015). Besides the possibility of an interaction of *helP* with its flanking T4P-system, there might be other mechanisms involved in how DEAD-box helicases could affect virulence. Tan et al. have reported on a DEAD-box helicase of *P. aeruginosa* that was essential for virulence and bacterial cytotoxicity in a mouse pneumonia model (Tan et al. 2016). Deletion of the DEAD-box helicase resulted in significantly lower expression levels of the T3SS effector protein ExoS and in a decreased production of proinflammatory cytokines and neutrophil infiltration in infected mice. However, we did not observe a differential expression of ExoS in *helP* positive isolates, but of ExoU levels, suggesting another potential linkage with the T3SS effectors.

Protein level analysis has also identified putative virulence and anti-virulence factors in *P. aeruginosa* bloodstream infection. Although strains were immediately conserved after detection and protein levels were determined in the first subculture after thawing, and therefore close to the conditions in the blood culture bottle, it is unknown how such protein level profiles would mirror pathogen protein levels in a patient’s bloodstream. Because of this limitation, we focused on *helP* as genomic biomarker due to its stability even under different pre-analytical conditions. We also hypothesized that protein level analysis can be a valuable tool in detecting additional virulence markers. Thus, we included factors arisen from this phenotype screening model into our final Cox regression prediction model.

High protein level of the flagellum basal body protein FliL was associated with increased mortality. The flagellum apparatus has been widely reported to be vital for virulence in *P. aeruginosa* (Kazmierczak et al. 2015). It is mainly important for swimming motility and attachment to host cells. Flagella components are known to bind to Toll-like receptor 5, thereby activating a mostly proinflammatory immune response (Zhang et al. 2003). During chronic infection in cystic fibrosis patients, flagellum expression is often downregulated to reduce inflammation (Mahenthiralingam et al. 1994). Our observation, together with the reported success of an anti-flagella vaccine in a clinical trial (Doring et al. 2007), makes the flagellum apparatus an interesting target for therapeutic virulence blockers in sepsis.

Another prognostic biomarker candidate is Prot214 which is annotated as bacterioferritin. Its role as risk factor for fatal outcome remains elusive. It is well known that an important line of defense against bacterial infection is the withholding of free iron since bacterial pathogens essentially depend on iron for replication and their pathogenic actions. In order to ensure sufficient iron levels in the bacterial cytosol and to also prevent iron-induced toxicity, cellular levels of free iron need to be highly regulated. In *P. aeruginosa*, two ferritin-like molecules are known to store iron intracellularly (bacterial ferritin A and bacterioferritin B) and both are considered obligatory for iron metabolism (Rivera 2017). These iron stores, suggested to be an important source for the heme prosthetic group of KatA, can increase resistance against hydrogen peroxide (Ma et al. 1999). This could be crucial for the rapid adaptation of invasive strains to new environments like the human blood and could augment pathogen survival against innate immune defense mechanisms. Nonetheless, the function of the bacterioferritin-like protein Prot214 as well as the anti-virulence capacity of the putative aminotransferase Prot330 during bacteraemia needs to be further investigated. This also applies to the discovery that GO-terms for peptidyl-histidine phosphorylation as part of two-component systems were enriched in the high risk accessory genome group (acc-cluster 2). The enrichment could reflect an improved ability for an immediate response to external stimuli. This could be advantageous in terms of a pronounced growth of invasive strains in different human body sites.

We conducted a systematic search for prognostic biomarker candidates in patients with *P. aeruginosa* bloodstream infection. Routine detection of highly virulent strains could result in administering high-dose combination therapy to those patients who need it most, providing a fair rationale for the additive toxicity. This is especially the case in *P. aeruginosa* bloodstream infection where combination therapy is thought to be superior over monotherapy, particularly when patients are at a higher risk of fatal outcome (Safdar et al. 2004; Park et al. 2012; Kim et al. 2014). Beside therapeutic management, detection of high-risk strains could also be followed by infection control measures like contact isolation. This would allow a reduction in the spread of virulent strains, an objective that is neglected by current infection control guidelines that tend to focus solely on a strain’s antibiotic susceptibility profile. Such practices could help in significantly reducing the more than 40,000 annual sepsis deaths alone in Europe and the hundred thousands more throughout the world. Our multi-omics approach has produced genomic and proteomic data identifying pathogen-derived prognostic biomarker candidates that are interwoven with treatment- and patient-related risk factors in a complex interplay. Our results reveal the importance of multi-omics approaches, which allow us to investigate multiple pathogen and host factors at the same time. Future studies that aim to validate these findings and to move confirmed pathogen-derived prognostic biomarkers into clinics are warranted.

## Methods

### Setting

We conducted a multicenter genomic cohort study in a 1500-bed tertiary teaching hospital, a 500-bed district hospital, and a 300-bed trauma center in Tübingen, Germany, and the surrounding community. The broad spectrum of medical services provided by these hospitals includes multiple medical and surgical specialties, pediatric units, dialysis and a maternity ward. Organ transplantations are carried out at the tertiary teaching hospital. The study is reported pursuant to the Transparent Reporting of a multivariable prediction model for Individual Prognosis Or Diagnosis (TRIPOD) and Strengthening the reporting of Genetic Risk Prediction Studies (GRIPS) statements (Janssens et al. 2011; Collins et al. 2015). Our study was approved by the local research ethics committee of the University of Tübingen (reference number: 364/2013R).

### Study design, patients and definitions

Adult patients (≥ 18 years) admitted to one of the participating hospitals were considered eligible when they were suffering from a blood stream infection (BSI) with ≥ 1 blood culture positive for *P. aeruginosa*. Patients were included once at the time of the first positive blood culture (index culture). Thirty-day mortality for any cause was the clinical endpoint while patient- and pathogen-related factors were regarded potential predictors of outcome.

Relevant patient data variables were defined as follows: site of infection (primary, secondary, vascular catheter-related) as classified by the International Sepsis Forum (Calandra and Cohen 2005); Charlson comorbidity score (Charlson et al. 1987); nosocomial infection (any infection ≥ 48 hours after hospital admission); immunosuppression (HIV and/or neutropenia with a neutrophil count ≤ 1000 cells/μl, and/or immunosuppressive chemotherapy within the previous two months and/or receipt of prednisolone ≥ 10 mg/daily or equivalent steroid dose). The physiological patient condition was determined using the simplified acute physiology score II (SAPS II) at the index culture day (Le Gall et al. 1993). A systemic administration of at least one antimicrobial agent to which the isolate was susceptible *in vitro* was defined an appropriate antimicrobial treatment.

### Species identification and susceptibility testing

Species identification was carried out using MALDI-TOF mass spectrometry and the Vitek 2 system (bioMérieux, Marcy l’Etoile, France). Minimum inhibitory concentrations were assessed by antibiotic gradient strips (MIC Test Strip, Liofilchem, Italy) and interpreted according to EUCAST breakpoints (version 8.0, 2018).

### Genomic data acquisition and analysis

Genomic DNA of *P. aeruginosa* has been sheared into 450 bp fragments using a focus-ultrasonicator (Covaris, Woburn, USA). Preparation of DNA libraries was done with the NEXTflex™ DNA Sequencing Kit (Bioo Scientific, Austin, USA) followed by sequencing at 2 × 125 bp on a HiSeq2500 platform (Illumina, San Diego, USA). SPAdes (version 3.7.0) has been selected as *de novo* assembly tool and assembled scaffolds were annotated using Prokka (version 1.11) (Bankevich et al. 2012; Seemann 2014). A gene presence-absence-matrix of all study isolates has been generated by Roary (version 1.006924) using a 95% minimum percentage identity for blastp (Page et al. 2015). Structure of the accessory genome was assessed using the Ward cluster algorithm with the Pearson similarity index (Stata version 12.1, Stat Corp., College Station, USA). Clustering was visualized using Biolayout (Theocharidis et al. 2009). Core genome construction was performed with Spine (version 0.1.2), and SNP were called using samtools (version 0.1.19) and GATK tools (version 3.2-2) (Li et al. 2009; Van der Auwera et al. 2013). The core genome maximum-likelihood phylogeny was generated using Gubbins (version 2.1.0) to account for genomic regions which had undergone homologous recombination (Croucher et al. 2015). A maximum of 10 iterations were used and a generalized time reversible (GTR) substitution model with a gamma distribution of rates. Gene ontology (GO) term enrichment analysis was conducted with Blast2GO (version 4.0.7) after genes from each group were clustered using CD-HIT-EST (version 4.6) with a 90% similarity threshold to remove redundancies (Conesa et al. 2005; Fu et al. 2012). The analysis was performed using a two-tailed Fisher’s exact test. The maximum false discovery rate (FDR) was set to 0.05 for reporting significant GO-terms and the Benjamini-Hochberg correction was used. Genomes from five strains were further determined on a PacBio *RS* II instrument. Each strain was sequenced in one SMRT cell resulting in coverage rates between 9-fold and 149-fold. Assembly of PacBio long reads was done using Canu version 1.5 (Koren et al. 2017), and contigs were subsequently polished with Pilon (version 1.22) to improve accuracy (Walker et al. 2014). For ID50, due to the low coverage of 9-fold, we used the SPAdes assembler version 3.9.0 (Bankevich et al. 2012) with Illumina short reads and the -- pacbio option. With this hybrid approach, we improved N50 statistic from 255,540 bp to 634,760 bp in this strain.

### Proteomic data acquisition and analysis

*P. aeruginosa* strains were grown overnight, and proteins were extracted as described elsewhere (Krug et al. 2013). Protein extracts were precipitated overnight with acetone and approximately 10 μg were loaded onto a NuPAGE Bis-Tris 4-12 % gradient gel (Thermo Fisher Scientific, Waltham, USA). Samples were let run approximately 10 mm into the gel and cut out as a single slice. In-gel digestion and peptide extraction were performed essentially as described previously (Borchert et al. 2010). Peptides were desalted using C18 StageTips (Rappsilber et al. 2007). LC-MS/MS analyses were performed on an EasyLC II nano-HPLC coupled to an LTQ Orbitrap Elite mass spectrometer (Thermo Fisher Scientific, Waltham, USA). LTQ Orbitrap Elite was operated in the positive ion mode. Samples were randomized before injection and a custom-made standard was measured in regular intervals to assess long-term performance of the MS.

Acquired MS spectra were processed with MaxQuant software package (version 1.5.2.8), with integrated Andromeda search engine (Cox and Mann 2008; Cox et al. 2011). Database search was performed against a *P. aeruginosa* database obtained from Uniprot (all strains), containing 103,188 protein entries, together with the custom-made database containing 30 additional entries which were not represented in the main database. Trypsin (full specificity) was set as the protease and the maximum number of missed cleavages was set to two. False discovery rate of 0.01 was set at the peptide and protein level. The label-free algorithm was enabled and a minimum of two unmodified peptide counts were required for quantification. Core proteome architecture was explored by using the Ward cluster algorithm with the correlation coefficient index (Stata version 12.1, Stat Corp., College Station, USA)

### Statistical analysis for virulence candidate assessment

A multi-level Cox regression analysis was applied to study the association between exposure (patient characteristics, geno- and phenotype of the pathogen) and the study endpoint (30-day all-cause mortality). Prior to testing, a variance range was set for all binary variables of interest to reduce dimensionality. Only those variables with a frequency ≥ 10% and ≤ 90% were tested. The final Cox regression model was built in a stepwise procedure. First, patient characteristics were individually tested and any variable with a p-value of < 0.2 was included in a multivariate model, wherein only variables with a p-value of ≤ 0.05 were retained, generating the clinical Cox regression model. In a second step, pathogen-related features were individually incorporated into the clinical Cox regression model. This led to three different screening models which integrated each one of the following information: accessory genome information (accessory genome gene screening model), phenotypic properties (natural log-transformed LFQ intensities of the core proteome and MICs = phenotypic screening model) or information about SNPs in known virulence factors (variant screening model) (Table S5). For SNPs, linkage disequilibrium was assessed and an R^2^ > 0.8 led to grouping and testing of one representative SNP for each group.

Integration of pathogen-related variables from each of the three screening models into the final multivariate model had to run through two selection processes: (i) Within each screening model, variables must have had a p < 0.05 and (ii) must have belonged to the 10% of variables with the lowest p-value from that model. In the final Cox regression prediction model, variables with a p ≤ 0.05 were retained. Hypothesis testing was performed by using the likelihood ratio test. The proportional hazards assumption was verified on the basis of Schoenfeld residuals. The final Cox regression prediction model was internally validated by bootstrapping (10000 replicates) and the jackknife method. Computations were done using Stata version 12.1 (Stat Corp., College Station, USA).

### Machine learning estimator search and optimization

The following were submitted for further evaluation using different machine learning approaches: datasets containing all features of the three screening models, a dataset containing the clinical risk factors and all pathogenic factors from the screening models (ALL) and the dataset with the variables from the final Cox regression prediction model (Final). For each algorithm, 30-day mortality was the outcome variable, and a model’s ability to predict the risk of a fatal case was assessed through a receiver operating characteristics analysis (ROC) measuring the area under the ROC curve (AUC) and through Matthews correlation coefficient. The scikit-learn toolbox version 0.19.1 was used for all calculations (http://scikit-learn.org/stable/). The following classification algorithms were tested on each dataset: random forest, support vector classifier, linear support vector classifier, k-nearest neighbour, and multi-layer perceptron.

The best estimator was searched on a training dataset that was comprised of 80% of the whole dataset. On each training dataset, we performed (i) no feature modification, (ii) dimensionality reduction using a principle component analysis with a maximum of 100 components and (iii) feature selection of the 5% features with the lowest univariate p-values. An exception was the dataset with the features from the final Cox regression prediction model where all calculations were only performed on the unaltered set of features. Hyperparameter tuning was conducted using the exhaustive grid search function (GridSearchCV), and estimator performance was evaluated by a ten-fold cross-validation. Here, the training set was split into 10 smaller sets. Subsequently, a model was trained on 9 folds of the training data and validated on the remaining part. The reported performance was the average of values computed in the loop. Potential over- and underfitting was determined by learning curves of the training datasets. Best estimators were finally evaluated on a hold-out dataset, which contained data the estimator has not seen before (remaining 20% of the data). This is to assess the likely “real-world” performance of the model estimator.

## Declarations

### Acknowledgements

We thank the directors, physicians, laboratory and nursing staff of the medical wards in all participating hospitals. We also extend our gratitude to Kerstin Fischer, Nadine Hoffmann, Sara Riedel-Christ, Silke Wahl, and Irina Droste-Borel for their excellent technical support. Our work was partially funded by the AKF fund of the University Hospital Tübingen (project number: E.03.43003) and the German Center for Infection Research (project number: TTU 08.702). The funders had no role in study design, data collection and analysis, in decision to publish, or preparation of the manuscript.

